# Use of an enclosed elk population to assess two non-invasive methods for estimating population size

**DOI:** 10.1101/2021.05.21.445203

**Authors:** Jennifer L. Brazeal, Benjamin N. Sacks

**Affiliations:** Mammalian Ecology and Conservation Unit, Veterinary Genetics Laboratory, School of Veterinary Medicine, University of California, Davis; Department of Population Health and Reproduction, School of Veterinary Medicine, University of California, Davis

## Abstract

Non-invasive genetic sampling and spatially explicit capture-recapture (SCR) models are used increasingly to estimate abundance of wildlife populations, but have not been adequately tested on gregarious animals such as elk (*Cervus canadensis*), for which correlated space use and movements violate model assumptions of independence. To evaluate the robustness and accuracy of SCR, and to assess the utility of an alternative non-invasive method for estimating density of gregarious ungulates, we utilized a tule elk (*Cervus canadensis nannodes*) population of known size within a fenced enclosure on the San Luis National Wildlife Refuge in central California. We evaluated fecal genetic SCR to camera trap-based random encounter model (REM) approaches to density estimation based on comparison to the true abundance. We also subsampled the dataset to explore the effects of varying search effort and elk density on the precision and accuracy of results. We found that SCR outperformed REM methods in the full datasets, and reliably provided accurate (relative bias <10%) and reasonably precise (relative standard error ≤20%) estimates of density at moderately low to high densities (6–17 elk/km^2^), when the subsampling scenarios yielded a minimum average of 20 recaptures. We also found that the number of samples used to construct detection histories was a reliable predictor of precision, and could be used to establish minimum sampling requirements in future population surveys of elk. Although field-testing in free-ranging populations is needed, our results suggest that non-invasive genetic SCR is a promising tool for future population studies and monitoring of elk and potentially other gregarious ungulates. In contrast, the REM estimate of density was highly inaccurate, imprecise, and highly sensitive to camera parameters.

## Introduction

Accurate estimates of abundance are of fundamental importance to the conservation and management of wildlife populations (Hauser et al. 2006, Engeman 2003, Nuno et al. 2015). Maintaining a sustainable population and setting well-informed harvest quotas for game species, such as elk (*Cervus canadensis*), are primary objectives of wildlife management agencies. Wildlife managers have traditionally monitored large ungulate populations using indices of abundance, such as pellet counts and hunter surveys, and counts or distance sampling from aerial surveys (Cogan and Diefenbach 1998, Jung and Kukka 2016, Terletzky and Koons 2016). However, indices are affected by observer, spatial, and temporal biases, while aerial distance sampling suffers from visibility bias, and is especially poorly suited to habitats with extensive canopy cover (Pollock and Kendall 1987, Amos et. al 2014).

In the past decade, the development and use of spatial capture-recapture (SCR) models to estimate population density and abundance has grown (Borchers and Fewster 2016), and non-invasive sampling methods that allow for individual identification can be used in conjunction with SCR to estimate population density for species that are wide-ranging and elusive (Kéry et al. 2010, Sollmann et al. 2011). Such sampling methods include the collection of genetic material left in the environment (e.g., hair, feces, feathers, etc.), as well as the collection of images from trail cameras for species with individually unique markings. Non-invasive SCR estimation of density has been used in a variety of species, including large carnivore species (e.g., Stansbury et al. 2014, Morehouse et al. 2016, Murphy et al. 2016), mesocarnivores (e.g., Monterroso et al. 2014, Thornton and Pekins 2015, Morin et al. 2016), ungulates (e.g., Brazeal et al. 2017), and other mammals (e.g., Moore and Vigilant 2014, Gerber and Parmenter 2015).

As an alternative to the capture-recapture modeling framework, Rowcliffe et al. (2008) developed a random encounter model (REM), designed to estimate density without the need for individual identification, and tested it empirically on a number of species, including various ungulates. The REM model assumes animals move across space in a random manner akin to gas particles, and uses encounter rates (i.e., photographic capture rates), the speed of animals, and the area over which the camera detects animals, to estimate density. This model has previously yielded estimates of density comparable to results from other methods in ungulate-focused studies, including zebra (*Equus Grevyi*), wild pig (*Sus scrofa*), moose (*Alces alces*), and roe deer (*Capreolus capreolus*) (Zero et al. 2013, Chauvenet et al. 2017, Pfeffer et al. 2018).

A potential challenge in estimating abundance and density of gregarious ungulates, such as elk, using SCR and REM arises from their tendency to move in groups. Most SCR models assume that animal activity centers (i.e., the centers of activity of individual animals during the sampling period), as well as our detections of individual animals, are independent of each other. There have been some exceptions in the literature to this standard. For example, Reich and Gardner (2014) developed an alternative point process model of animal activity centers for territorial animals. In addition, for group-living species, we can obtain an estimate of group density from SCR by using a group as a sampling unit rather than an individual. Combined with an independent estimate of average group size, group density can be used to obtain individual animal density (e.g., Russell et al. 2012). When applied to natural populations, in both cases, modeling the distribution of individual activity centers as independent yielded similar density estimates as models that accounted for dependence among locations, suggesting that SCR models may be robust to violations of the independence assumption. The REM model also assumes independence in encounter rates of individuals, and the recommended solution for violations of this assumption in group species is to estimate group density and size (Rowcliffe et al.2008, Zero et al. 2013), then derive individual density, as in SCR.

Our primary objective was to test and compare noninvasive genetic SCR and camera REM methods in terms of accuracy and precision of density estimates for a group-living species using data gathered from an enclosed population of tule elk (*C. c. nannodes*) of known size. We also used random subsampling from a dataset that included detections from all individuals within the enclosure to evaluate the performance of SCR under varying levels of sampling effort and low to high densities of elk. Lastly, we used the results from subsampling to identify a sampling effort variable that could be used to predict precision in density estimates from surveys using non-invasive fecal genetic sampling and SCR methods.

## Study Area

The San Luis National Wildlife Refuge spanned 108.46 km^2^ in the San Joaquin Valley near the city of Los Banos in Merced County, CA. Dominant habitat types across the Refuge included freshwater wetlands, valley riparian, annual grasslands, and oak woodlands. The study area encompassed a 49 km^2^ enclosed area that was predominately annual grasslands and riparian areas, dominated by willow (*Salix* spp.) and cottonwood (*Populus fremontii*). The climate was Mediterranean, with arid, hot summers and wet, mild winters. Mean daily temperature during the study period was 24 °C with a minimum of 8.9 °C and a maximum of 37 °C (California Department of Water Resources, Station ID: SAN, Jul-Sep, 2016). A small population of tule elk (historically ranging from 18 – 73 individuals) was confined within the fenced enclosure since 1974 to serve as a source for periodic translocations and reintroductions to former range and for conducting applied research. At the time of our study, the enclosure contained 72 individuals (26 ad F, 35 ad M, and 11 juv) based on a census count conducted on July 27, 2016, shortly after the start of our study (14 July 2016).

## Methods

### Spatial Capture-Recapture

#### Sampling

To ensure an approximately even distribution of sampling, we overlaid a 2.25 km × 2.75 km rectangular polygon across the enclosure boundary, which we divided into 250 × 250 m plots. For the 56 plots overlapping the enclosure boundary with more than 50% of their area, we attempted to sample, at a minimum, the vertical length of each plot, traversing it from the north to the south edges. We recorded and saved our walking paths in handheld GPS units so we could use the length of the distance walked within a grid cell as a measure of spatial search effort. We sampled during summer, from 14 July to 1 September 2016. We selected this time period to avoid the rutting and calving periods, as both could systematically alter home range sizes and group membership, as well as violate the assumption of demographic closure.

To collect samples, we walked while scanning the ground for pellets 1 m on either side of our walking path. When we encountered an elk pellet group, we collected pellets that appeared to <2 weeks old (i.e., no surface cracks). From each distinct pellet group, we collected 3–6 pellets in a 50-mL conical vial and stored them in 95–100% ethanol. We recorded the GPS coordinates of each sample collection. Because SCR models are capable of estimating the probability of detection using information solely from spatial “recaptures,” we did not systematically resample grid cells, though we sometimes resampled a grid cell along an alternative path to increase sample size.

#### DNA analysis

We completed all laboratory analyses at the Mammalian Ecology and Conservation Unit of the University of California Davis Veterinary Genetics Laboratory. To extract DNA from the epithelial cells on the surface of the pellets, we followed a modified version of the protocol described in Brazeal at al. (2017) for deer pellets, to accommodate the larger size of the elk pellets. Specifically, we used only 1–2 elk pellets, and a larger volume (2.5 mL instead of 1 mL) of buffer ATL (Qiagen, Valencia, CA, USA) to wash the epithelial cells from the pellet surface. We then extracted DNA from the resulting suspension of epithelial cells using the DNeasy 96 Blood and Tissue Kit (Qiagen, Valencia, CA), according to the manufacturer’s protocol. For each 96-sample extraction set, we included a negative control to detect any contamination during the extraction process. We attempted to amplify via polymerase chain reaction (PCR) each DNA sample in two independent PCR reactions to reduce error from random allelic dropouts. In each PCR reaction set, we included two negative PCR controls to detect possible contamination. We genotyped the DNA samples using a single multiplex assay designed to be sufficiently polymorphic for tule elk (Sacks et al. 2016). We included 11 microsatellite markers: TE179, TE159, TE85, TE132, TE84, TE185, TE45, TE182 (Sacks et al. 2016), T26, T193 (Jones et al. 2002), and a sex typing marker from the Y chromosome (SRY; Wilson and White 1998). We used an ABI 3730 (Applied Biosystems, Grand Island, NY) and internal size standards (500-LIZ; Applied Biosystems) for electrophoresis, and manually scored alleles using electropherograms in Program STRand (version 2.4.89; Toonen and Hughes 2001).

We combined replicate multi-locus genotypes into single composite consensus genotypes for each sample, and excluded consensus genotypes that did not have 100% amplification success across loci from downstream analyses. We used R package allelematch to assign consensus genotypes to individual clusters, based on an optimal threshold of mismatching loci (Galpern et al. 2012). We also calculated the unbiased probability of identity (i.e., probability of multi-locus genotypes matching from any 2 random individuals, P_ID_) and the more conservative probability of identity for siblings (P_ID_Sib) (Waits et al. 2001).

### Density estimation

We implemented SCR models specific to data collected from area searches in a maximum likelihood framework (Efford 2011) in R (R Version 3.5.1, www.R-project.org, accessed 25 September, 2018) using the package SECR (Efford 2018). We assumed a uniform and independent distribution of activity centers (i.e., the center of an animal’s space use activity during the sampling period, which we hereafter refer to as its “home range”). Because the elk were enclosed by an impenetrable fence barrier, we parameterized the model using the enclosure boundary as the habitat mask (i.e., the area that includes the home range centers of all animals exposed to sampling over the course of the study). We used the extent of the enclosure boundary to delineate a rectangular polygon as the search area. We used the hazard exponential function to describe the relationship between the expected number of detections *λ(d)* and the geometric overlap between the animal’s home range center and the search area:

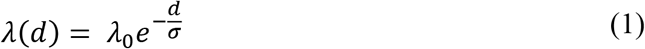

where *d* is the distance between the animal’s home range center and where it was detected, *σ* is a scaling parameter that describes the rate of decline in detection as the distance between the animal’s home range center and its detection location increases, and *λ*_0_ is the intercept of the detection function. The overall expected encounter rate of an animal with a home range centered on *x* would then be based on the overlap between the radially symmetric and circular probability density of the animal’s home range and the search area polygon, and was estimated using the following equation (Efford 2018):

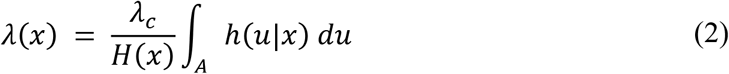

In the equation above, ℎ(*u|x*) is the hazard exponential function described in eq. 1, integrated over the search area polygon (A), *λ_c_* is the expected number of detections of an animal with a home range enclosed entirely within the search area, and *H*(*x*) is the cumulative detection function, integrated over the state space (i.e., the enclosure boundary), included as a normalizing constant. The expected number of detections (*λ*(*x*)) for an animal increases as the areal overlap of its home range with the search area polygon increases. The probability of detecting an animal (*g*(*x*)) is also higher if more of its home range overlaps the search area, and is related to the expected encounter rate with the following equation:

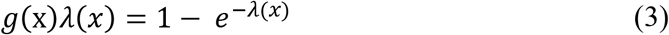

We estimated density (*D*) in a series of models that we parameterized based on expected sources of heterogeneity in the detection parameters. Because we expected the scale of movement (*σ*), and possibly the expected number of detections (*λ_c_*), to vary by sex, we allowed *σ* and *λ_c_* to vary using finite mixture models for individual heterogeneity (Pledger 2000). Our first set of models (s0–s2) considered sex to be a known 2-class mixture factor. Since all classes (i.e., sexes) were known, this model was equivalent to a model using sex as a grouping factor, but also allowed estimation of the sex proportions from an additional parameter, *pmix* (Borchers and Efford 2008). In s0, our null model for sex, the detection parameters (*σ* and *λ_c_*) were constant, while in s1, we varied *σ* by sex, holding *λ_c_* constant, and in s2, both *σ* and *λ_c_* varied by sex. We expected space use to vary by sex only in adult elk, but because we had no way to assign age classes to individual identities, differences by sex might have been obscured by similarities in space use among calves and yearlings. As such, we considered an alternative set of models, in which we modelled individual heterogeneity by finite latent 2-class mixture models (h2a–h2c) (Pledger 2000, Borchers and Efford 2008). These models attempt to capture individual heterogeneity in detection probabilities by estimating different detection probabilities for 2 or more latent classes of unknown identity. Mirroring the first set, h2a assumed that *σ*, and *λ_c_* were constant, while h2b varied only *σ* between the latent classes and h2c assumed that both *σ* and *λ_c_* varied between classes. We used Akaike’s Information Criterion (AIC) to rank the models within each set (Burnham and Anderson 2002).

To explore the effect of sampling effort on the accuracy of SCR results (i.e., evaluated against the known number, 72 individuals), we subsampled from our empirical noninvasive genetics dataset by randomly selecting different numbers of 250 × 250 m plots and different numbers of elk individuals, and estimated *D* from the reduced datasets. For each subsampled dataset, the reduced sampling layouts and reduced elk populations were subsampled separately, then combined into one dataset by removing any detections that fell outside the subsampled plots. First, we randomly selected *j* plots (*j* = 10, 15, 20, 25, 30, and 35 plots) within the enclosure to simulate different sampling layouts. For each reduced plot scenario, we replicated sampling 10 times with replacement, for a total of 60 reduced sampling layouts (e.g., 10 random sets of 10 plots, 10 random sets of 15 plots, etc.). Based on the census count, our survey recorded all individuals within the enclosure, and we were, therefore, also able to explore the effect of varying elk density on the accuracy of SCR results by subsampling. Specifically, we randomly selected multiple subsets of *k* individuals (*k* = 20, 30, 40, 50, and 60 elk) and replicated each reduced elk density scenario 10 times. Observed densities of wild elk populations in North America can vary widely, from <1 elk/km^2^ to >30 elk/km^2^, and our reduced density scenarios corresponded to a range of moderately low to high densities of elk (6–17 elk/km^2^) (Stewart et al. 2009, Proffitt et al. 2015). Our subsampling scenarios resulted in search efforts ranging from 1.0–6.6 km searched for each km^2^ of the study area (km/km^2^), which was a higher intensity of effort compared to previous studies using non-invasive genetic methods to sample wild ungulate populations for SCR. For example, prior studies on mule deer (*Odocoileus hemionus*) in California used sampling efforts ranging from 0.01–0.96 km/km^2^ to estimate density (Brazeal et al. 2017, Furnas et al. 2018). However, test scenarios using comparable levels of sampling consistently resulted in <20 recaptures, perhaps because of differences in the scale of movements of animals in our enclosure compared to wild ungulates, or because we focused on spatial allocation of effort rather than resampling the same areas.

Subsampling resulted in 3,000 reduced-capture history datasets (i.e., 6 plot scenarios × 10 replicates × 5 elk density scenarios × 10 replicates). For each of the 3,000 subsampled datasets, we estimated *D* using the model found to be most parsimonious based on AIC (see Results). We derived the estimated population size 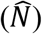 as 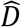 multiplied by 3.49 km^2^, the area inside the fenced enclosure (Efford and Fewster 2013). We used the relative standard error 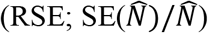 to evaluate the precision of the models under the different subsampling scenarios and the difference between the average 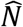 under each subsampling scenario and the subsampled number of elk (*N*) to estimate relative bias 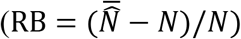

We were also interested in determining a useful predictor for the RSE of 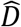 that managers could use as a threshold parameter to set sampling goals for monitoring surveys, based on a desired level of precision in 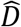. We assumed that the RSE of 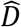 would have a negative relationship with both search effort and the subsampled density of elk. In other words, as search effort and the density of elk on the landscape increased, we expected a decrease in the RSE of 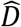 (i.e., an increase in precision). For each dataset, we converted the simulated population size to a density of elk by dividing *k* subsampled individuals by the area of the enclosure (3.49 km^2^). We calculated the search effort for each subsample layout from the sum of the total length (km) of GPS tracks recorded in *j* plots. We also expected that RSE of 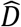 would decrease with the number of genotyped pellets used to construct encounter histories of individuals, and tested models with the number of pellets used in a given subsampled dataset as the predictor for RSE. As the total number of pellets used for a given subsampled dataset was a function of both the search effort and the density of elk in the area, it would be a more efficient predictor of RSE if the prediction error from the model was low.

For both sets of models, we performed generalized linear models, and considered both the gamma and inverse Gaussian probability distributions for the error in RSE. These probability distributions were both suitable for modeling continuous, non-negative response variables, where the conditional variance of the variable increases with its expected value (Fox 2008), as we expected of the RSE of 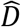. We used AIC to compare the model likelihoods. We also used the boot package, ver. 1.3-19 in R to perform 10-fold cross-validation (CV) in order to estimate and compare prediction error of the models (Davison and Hinkley 1997, Canty and Ripley 2017). In the models with both search effort and elk density as predictor variables, we tested for multicollinearity using a Fararr test (Farrar and Glauber 1967).

### Random Encounter Model

#### Sampling

For the camera portion of the study, we systematically placed cameras at three central locations within the enclosure. We used Cuddeback Long Range IR model E2 cameras. We divided the 3.49 km^2^ enclosure into three irregular polygons, each approximately 1.2–1.3 km^2^. At the centroid of each polygon, we designated three camera locations, situated 100 m apart in a triangular configuration, and placed north-facing and south-facing cameras at each location, resulting in a total of 18 cameras at 9 camera locations (Fig. 1). We wanted to effectively sample the entire enclosure, and expected that placing clusters of cameras in approximately 1-km^2^ areas would adequately capture the majority of the elk within the enclosure. Further, we expected elk movement to be random with respect to the camera locations, as we did not consider elk distribution or movement in our placement of cameras. We placed the cameras on trees, when available, approximately 1.25 m above the ground, but for 4 locations, we installed metal posts for camera placement, and at 2 locations, we attached cameras to telephone poles. We set the delay interval between photos to be 30 s. We switched out the SD memory cards in each camera weekly. The cameras were active from 13 July, 2016 to 30 August, 2016 (49 days), but for analysis we only used photos captured between 20 July, 2016 to 30 August, 2016 (42 days). We truncated the sampling period to allow elk to acclimate to the presence of the cameras before using encounter rates in the REM model.

**Figure 1.**
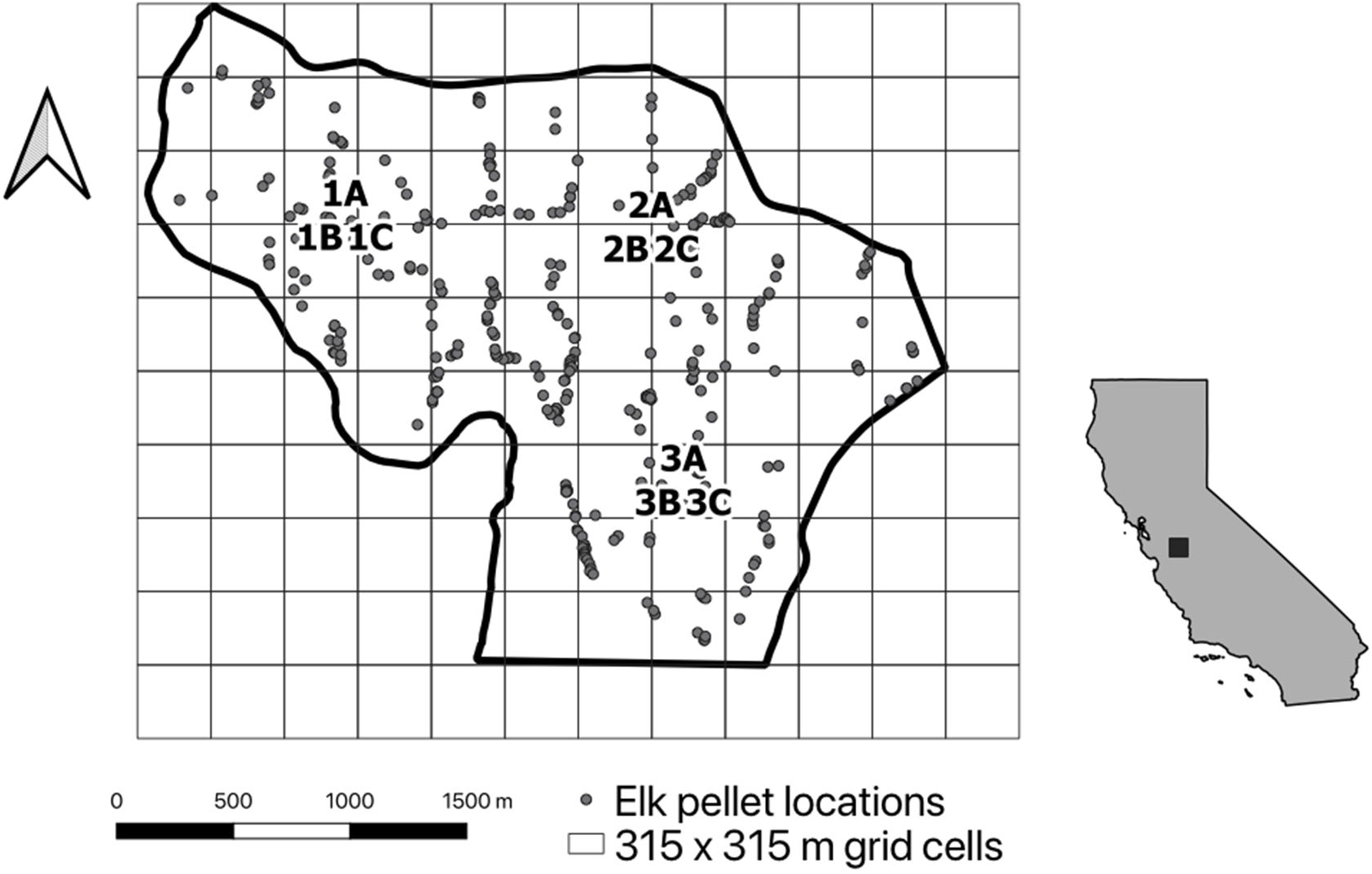
Map showing the boundary of the elk enclosure at the San Luis National Wildlife Refuge, near Los Banos, CA. Points show the locations of elk pellet samples collected between 14 July and 1 September of 2016 for use in genetic spatial capture-recapture analysis. At each of 9 camera locations (1A-3C), a north and south-facing camera was placed, for a total of 18 cameras. We collected photos of elk between 20 July and 30 Aug 2016.

#### Density estimation

The random encounter model required estimation of 4 parameters to estimate *D*. These include the encounter rate of animals (*y/t* = count of photos/time), animal velocity (*v*), the radius of camera detection (*r*), and the angle of camera detection (*θ* radians). For animals that move in static groups, group 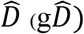 should be obtained instead by estimating the encounter rates of groups, and an independent estimate of group size (*g*) can be used to convert 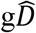 to 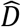 (Rowcliffe et al. 2008, Zero et al. 2013). The REM model modifies the ideal gas model to estimate *D* using the following equation:

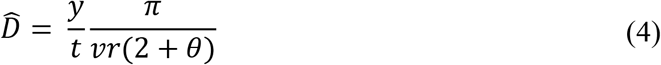

Because elk move in groups, we estimated group *y*/*t* for each camera by counting each independent photo capture of elk as a single encounter, regardless of the number of elk in the photo. If a camera captured the same group in consecutive photos, only the first photo was counted. We took into account both the time in between successive photo captures and the group composition when eliminating duplicate captures. We estimated *v* using GPS collar data that recorded locations every hour between 16 November, 2016 and 9 December, 2016 for 2 elk in the enclosure (1 adult male and 1 calf male). We selected an adult male to represent the movement rates of bull elk within the enclosure, and a calf to represent the movements of both cows and calves. We obtained an estimate of *v* in km/day by summing the total distance moved by both individuals over each full day they were monitored and dividing this distance by the total number of animal-days monitored. For an estimate of average *g*, we opportunistically counted and recorded the number of elk per grouping we sighted on sampling days and because the distribution of counts was right-skewed, we back-transformed the average of the log-transformed counts. We estimated camera detection parameters (*r* and *θ*) by marking distances from 5–18 m in a straight line from the center of the camera lens (line *A*) and every 0.5 m up to 5 m on a line perpendicular to the camera line of sight (line *B*), at a distance of 5 m from the camera. We approached the camera perpendicular to the line of sight at varying speeds and distances, and examined the resulting photos to record the distances at which the camera triggered. We calculated *θ* using the equation 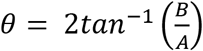 and *r* from the equation 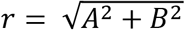 on the geometry of right triangles. As in the original version of the model, we assumed that the variance of camera detection parameters was 0, but in addition to measurement error, there can be variation in triggering rate due to factors such as the amount of available sunlight, the temperature, and the size of animals (Rowcliffe et al. 2011, Wellington et al. 2014). To examine how sensitive results from REM were to error in the estimation of these parameters, we conducted a sensitivity analysis to assess the changes in 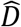 as *r* and *θ* varied. Following Rowcliffe et al. (2008), we calculated the variance of *y*/*t* by bootstrapping with replacement 10,000 times, with resampling stratified by camera location (i.e., the centroids of each of the 3 sample polygons), and the variance of 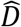 using the delta method (Seber 1982, Powell 2007).

## Results

### Spatial Capture Recapture

#### Genetic Samples

We sampled 57 250 × 250 m plots, and the average length of the GPS tracks that we recorded as we searched for pellets was 591.2 (SD = 440.7) m per plot, with lengths ranging from 13.74 m to 1,887 m. We collected a total of 483 pellet samples. Of the 483 samples, 326 amplified successfully at all 11 loci (67.5% of the samples), and we used these samples to determine individual identifications. We found 2 alleles to be the optimal allele mismatch threshold, meaning that individuals could be considered the same individual only if they matched at 21 or more of 23 allelic positions (11 loci and the sex marker, SRY). The overall P_ID_ was 2.6 × 10^−3^ and P_ID_Sib was 0.012 (Table 1). From the 326 samples, we detected and identified all 72 individuals in the enclosure. Individuals were recaptured 1–12 times, with an average of 4.6 (SD = 2.9) spatial captures per individual.

**Table 1.**
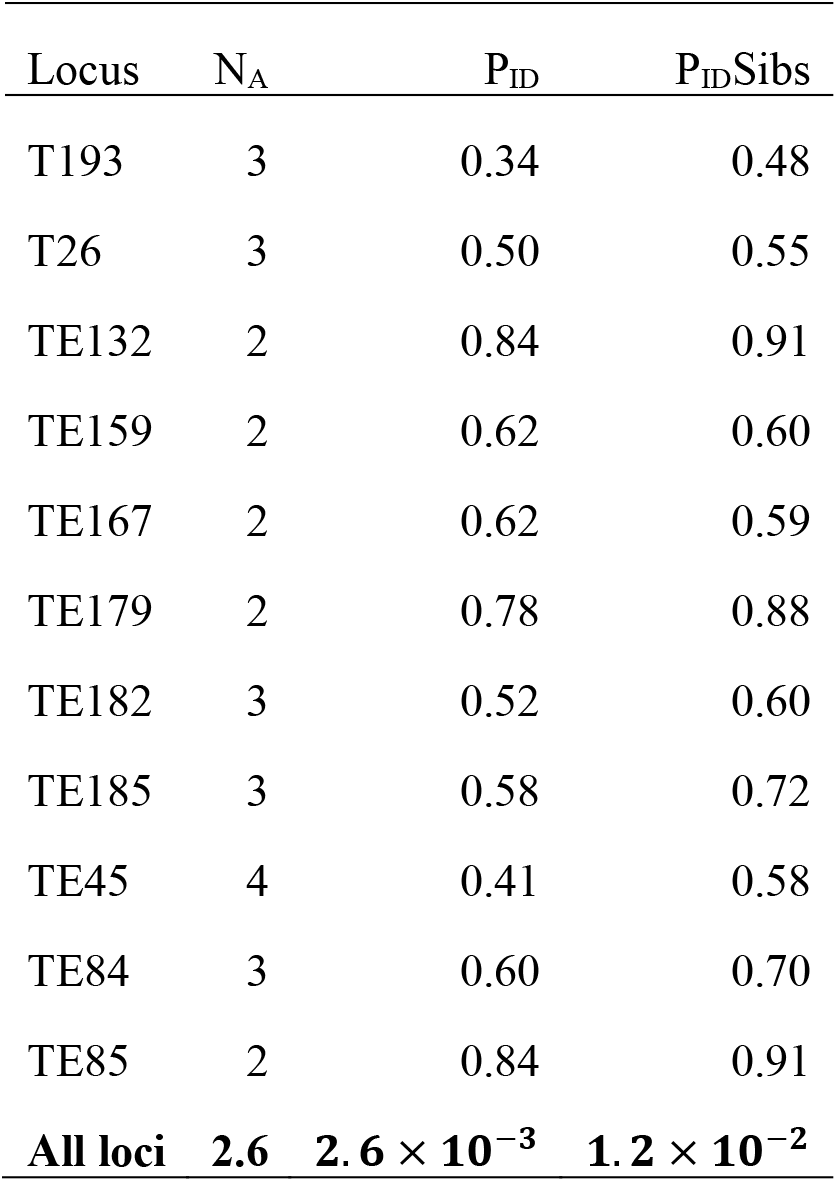
Estimates of allelic richness (N_A_), probability of identity (P_ID_), and probability of identity for siblings (P_ID_Sib) for each locus of the 11 loci used to identify individual tule elk from fecal samples collected from the San Luis National Wildlife Refuge in Los Banos, CA. The final row shows the average number of alleles per locus, and the P_ID_ and P_ID_Sib for the multilocus genotypes, obtained by multiplying the P_ID_ and P_ID_Sib estimates across all loci.

#### Density

For the models in which we used all data and sex was a 2-class mixture factor, the null model (s0) was the highest-ranking model by AIC (AIC weight = 0.64), though models s1 and s2 were within 4 ΔAIC of s0 (Table 2A). All three models had nearly identical results for 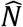, with point estimates rounding to 73 elk with high precision (RSE = 0.12). The model s0 estimate for *λ_c_* was 5.7 (SE = 0.37) expected detections and for *σ* was 334 (SE = 21) m. In both s1 and s2, the *pmix* ratio estimated the female proportion to be 0.37 (SE = 0.06) and the male proportion to be 0.63 (SE = 0.06).

**Table 2.** Model selection rankings and results from genetic spatial capture-recapture analysis using all elk pellet samples collected summer 2016 inside an elk enclosure at the San Luis National Wildlife Refuge in Los Banos, CA. We used Aikaike’s Information Criterion to rank the models (Burnham and Anderson 2004). Columns show the model name, the parameter structure (Parameters), the number of parameters used in the model (k), the AIC value, the delta AIC (ΔAIC) and AIC weight (AIC wt.). The estimated abundance of elk in the enclosure 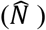 and standard error (SE) are also shown, as well as the relative standard error 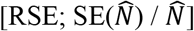. The *pmix* values show the estimated mixture proportion of each class. In table (A.), models s0-s2 used sex as a grouping factor to model the scale of movement of elk (sigma, *σ*) and the expected number of detections for an individual elk (λ_c_). In table (B.), models h2a-h2c assumed a finite latent 2 class mixture that we used to model *σ* and λ_c_ (Pledger 2000). We used the conditional likelihood to fit all models, and density was therefore a derived estimate using a Horvitz-Thompson-like estimator (Borchers and Efford 2008). The estimate of population size (N) and its SE was calculated by multiplying D by the area inside the elk enclosure (49 km^2^).

| Model | Parameters | k | AIC | ΔAIC | AIC wt. | $\hat{N}$ (SE) | RSE | 95% CI | F *pmix* (SE) | M *pmix* (SE) |
| --- | --- | --- | --- | --- | --- | --- | --- | --- | --- | --- |
| s0 | $\lambda_c \sim 1 \mid \sigma \sim 1$ | 3 | 10439.67 | 0.00 | 0.64 | 73.36 (8.65) | 0.12 | 58.27–92.37 | NA | NA |
| s1 | $\lambda_c \sim 1 \mid \sigma \sim \text{sex}$ | 4 | 10441.47 | 1.80 | 0.26 | 73.37 (8.65) | 0.12 | 58.27–92.37 | 0.37 (0.06) | 0.63 (0.06) |
| s2 | $\lambda_c \sim \text{sex} \mid \sigma \sim \text{sex}$ | 5 | 10443.24 | 3.58 | 0.11 | 73.37 (8.65) | 0.12 | 58.29–92.39 | 0.37 (0.06) | 0.63 (0.06) |

B.
| Model | Parameters | k | AIC | ΔAIC | AIC wt. | $\hat{N}$ (SE) | RSE (%) | 95% CI | h1 *pmix* (SE) | h2 *pmix* (SE) |
| --- | --- | --- | --- | --- | --- | --- | --- | --- | --- | --- |
| h2c | $\lambda_c \sim \text{h2} \mid \sigma \sim \text{h2}$ | 5 | 10326.32 | 0.00 | 0.99 | 88.56 (18.62) | 0.21 | 58.91–133.14 | 0.38 (0.10) | 0.62 (0.10) |
| h2b | $\lambda_c \sim 1 \mid \sigma \sim \text{h2}$ | 4 | 10335.20 | 8.89 | 0.01 | 90.34 (26.19) | 0.29 | 51.77–157.65 | 0.34 (0.16) | 0.66 (0.16) |
| h2a | $\lambda_c \sim 1 \mid \sigma \sim 1$ | 2 | 10342.40 | 16.08 | 0.00 | 73.36 (8.65) | 0.12 | 58.27–92.37 | NA | NA |

In contrast to models s0–s2, the models that assumed latent individual heterogeneity in the detection parameters (h2b and h2c) were ranked higher than the null model (h2a), but *D* estimates from these models were positively biased, with 95% confidence intervals that did not include the census count of 72 elk (Table 2B). The top ranking model by AIC was model h2c (AIC weight = 0.99), which varied *λ_c_* and *σ* by the 2 latent classes, while the null model (h2a) was the lowest ranking model (ΔAIC = 16.08). The positive bias in the estimate of *D* from model h2 likely arose from an inability to estimate accurately detection rates for both latent groups. The proportions between the sexes in model s2 and the latent groups in model h2 were similar, but the expected detection rates (*λ_c_*) for each group were quite different. From model s2, the male proportion of the population was estimated to be 0.63 (SE = 0.06), with the remaining proportion female, while the proportion of the larger latent group in model h2 was estimated to be 0.62 (SE = 0.10). Males from model s2 had an expected *λ_c_* of 5.5 (SE = 0.5) detections, which was similar to the female 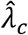 of 5.9 (SE = 0.6) detections. Meanwhile, model h2 estimated the larger latent group to have an expected *λ_c_* of 7.0 (SE = 0.10) detections, while the smaller group had an expected *λ_c_* of only 0.82 (SE = 0.87) detections. The high RSE of *λ_c_* for the smaller group in model h2 (RSE = 1.06) suggested a large amount of uncertainty in the estimate of *λ_c_*. The much lower estimated *λ_c_* for 38% (SE = 0.87%) of the population in model h2 likely resulted in the larger estimate of *D*.

Estimates based on application of the null model with sex as a 2-class mixture factor, subsampling the plots within the enclosure and simulating reductions in the elk density generally exhibited a slight negative bias, though in the majority of cases |RB| was <10% (Table 3). At the lowest sampling effort (15 plots), |RB| was >10% in some cases. In addition, when sampling only 15 plots at a true subsampled *N* of 10 or 20 elk, 3 models produced unreasonable estimates of 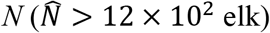 and were removed from the summary statistics. The RSE ranged from 0.14–0.54, decreasing with increases in both sampling effort and the simulated density of elk. For subsampling scenarios with an average minimum of 20 recaptured individuals, RSE was always ≤ 0.20, while |RB| was always <10% (Table 3).

**Table 3.** Average estimates of population size 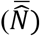, relative bias 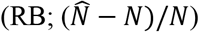, and average relative standard errors 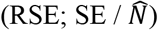 based on random subsamples from a full dataset of spatial capture-recapture detections for a known population of tule elk enclosed in a 3.49 km^2^ wildlife refuge in Los Banos, CA. We simulated random samples from the full empirical dataset 10 times for each number of 315 × 315 m grid cells (No. grids) within the enclosure. For each of the grid cell subsamples, we randomly subsampled the total number of elk (*N* = 72) 10 times to produce a new true population size (*N*) for a total of 100 replicates for each combination of No. grids and *N*. We calculated the average estimates of *N*, the RB and the RSE from the results for each subsampling scenario. The table also shows the average number of samples per replicate model 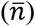 and the average number of recaptured individuals 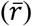.

| No. grids | $N$ | $\bar{N}$ | RB | RSE | $\bar{n}$ | $\bar{r}$ |
| --- | --- | --- | --- | --- | --- | --- |
| 15 | 10 | 9.48 | -0.05<sup>a</sup> | 0.54<sup>a</sup> | 11.95 | 3.36 |
| 20 | 10 | 9.47 | -0.05<sup>a</sup> | 0.43<sup>a</sup> | 16.58 | 4.42 |
| 25 | 10 | 9.25 | -0.08 | 0.40 | 21.45 | 5.46 |
| 30 | 10 | 9.29 | -0.07 | 0.37 | 24.73 | 5.96 |
| 35 | 10 | 9.61 | -0.04 | 0.35 | 30.01 | 6.74 |
| 15 | 20 | 19.97 | 0 | 0.39 | 21.60 | 6.04 |
| 20 | 20 | 19.64 | -0.02 | 0.30 | 30.72 | 8.11 |
| 25 | 20 | 18.81 | -0.06 | 0.27 | 39.96 | 10.47 |
| 30 | 20 | 19.11 | -0.04 | 0.26 | 45.71 | 11.73 |
| 35 | 20 | 19.34 | -0.03 | 0.24 | 55.64 | 13.52 |
| 15 | 30 | 28.08 | -0.06 | 0.28 | 33.61 | 9.24 |
| 20 | 30 | 28.90 | -0.04 | 0.23 | 48.59 | 13.22 |
| 25 | 30 | 27.93 | -0.07 | 0.21 | 62.58 | 16.27 |
| 30 | 30 | 28.70 | -0.04 | 0.20 | 72.73 | 18.63 |
| 35 | 30 | 29.10 | -0.03 | 0.20 | 86.39 | 20.96 |
| 15 | 40 | 35.19 | -0.12 | 0.24 | 43.70 | 11.92 |
| 20 | 40 | 37.07 | -0.07 | 0.20 | 62.13 | 16.37 |
| 25 | 40 | 36.26 | -0.09 | 0.19 | 81.43 | 20.65 |
| 30 | 40 | 37.45 | -0.06 | 0.18 | 92.15 | 22.94 |
| 35 | 40 | 38.34 | -0.04 | 0.17 | 112.41 | 26.42 |
| 15 | 50 | 42.95 | -0.14 | 0.21 | 56.98 | 15.68 |
| 20 | 50 | 46.02 | -0.08 | 0.17 | 82.42 | 21.71 |
| 25 | 50 | 45.33 | -0.09 | 0.16 | 107.09 | 26.82 |
| 30 | 50 | 46.85 | -0.06 | 0.16 | 120.99 | 29.73 |
| 35 | 50 | 47.98 | -0.04 | 0.15 | 146.34 | 33.96 |
| 15 | 60 | 51.98 | -0.13 | 0.19 | 66.1 | 18.10 |
| 20 | 60 | 55.42 | -0.08 | 0.16 | 95.28 | 25.34 |
| 25 | 60 | 54.42 | -0.09 | 0.15 | 123.18 | 31.11 |
| 30 | 60 | 56.11 | -0.06 | 0.14 | 140.78 | 35.04 |
| 35 | 60 | 57.17 | -0.05 | 0.14 | 171.18 | 40.22 |

<sup>a</sup>. We removed 3 replicates from the 15 grid cell/10 elk and the 20 grid cell/10 elk subsampled datasets because the $N$ estimates were much larger than true $N$ ( $>10^3$ times larger) and/or models failed to provide estimates of the SE. We did not include these results to estimate RB and RSE because these results would not typically be considered valid in an empirical dataset.

In terms of modeling precision (RSE) based on subsamples of the comprehensive dataset, the top-ranking model by AIC, with 100% of the AIC model weight, was the model that used number of samples as the predictor variable (Table 4). Modeling RSE by the number of samples using an inverse Gaussian distribution of errors also resulted in the lowest CV (1.41 × 10^−3^) out of all the candidate models. As the number of samples increased, the RSE decreased (β = 0.35, P < 0.001) (Fig. 2B). If the desired level of precision was, for example, an RSE ≤20%, the model predicted that at least 72 samples that successfully genotyped at all markers would need to be collected for this study, regardless of elk density or search effort. The observed number of samples that yielded *D* estimates with an RSE of 19.5–20.5% ranged from 56–100 pellets.

**Table 4.** Tables show the AIC model rankings (Burnham and Anderson 2004) and 10-fold cross validation error for generalized linear models that predict relative standard error 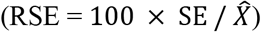 of results from genetic spatial capture-recapture, by km of search effort, density of elk (D), and the number of samples used in the analysis (No. samples). The data for the models were generated by random subsampling from the full dataset to reduce the number of 250 × 250 m grids sampled in the enclosure and the number of elk available for sampling.

| Model | GLM Family | Link function | k | AIC | $\Delta AIC$ | AIC wt. | Cross-validation error (K=10) |
| --- | --- | --- | --- | --- | --- | --- | --- |
| RSE ~ No. Samples | Inverse Gaussian | Inverse-squared | 3 | -15644.55 | 0.00 | 1.00 | 1.37E-03 |
| RSE ~ D + Search effort + D * Search effort | Inverse Gaussian | Inverse-squared | 5 | -14542.02 | 1102.52 | 0.00 | 3.08E-03 |
| RSE ~ D + Search effort | Inverse Gaussian | Inverse-squared | 4 | -13166.12 | 2478.43 | 0.00 | 3.96E-03 |
| RSE ~ D + Search effort + D * Search effort | Gamma | Inverse | 5 | -12801.81 | 2842.73 | 0.00 | 3.19E-03 |
| RSE ~ D + Search effort | Gamma | Inverse | 4 | -12659.78 | 2984.77 | 0.00 | 3.22E-03 |
| RSE ~ No. Samples | Gamma | Inverse | 3 | -12399.962 | 3244.58 | 0.00 | 3.02E-03 |

**Figure 2.**
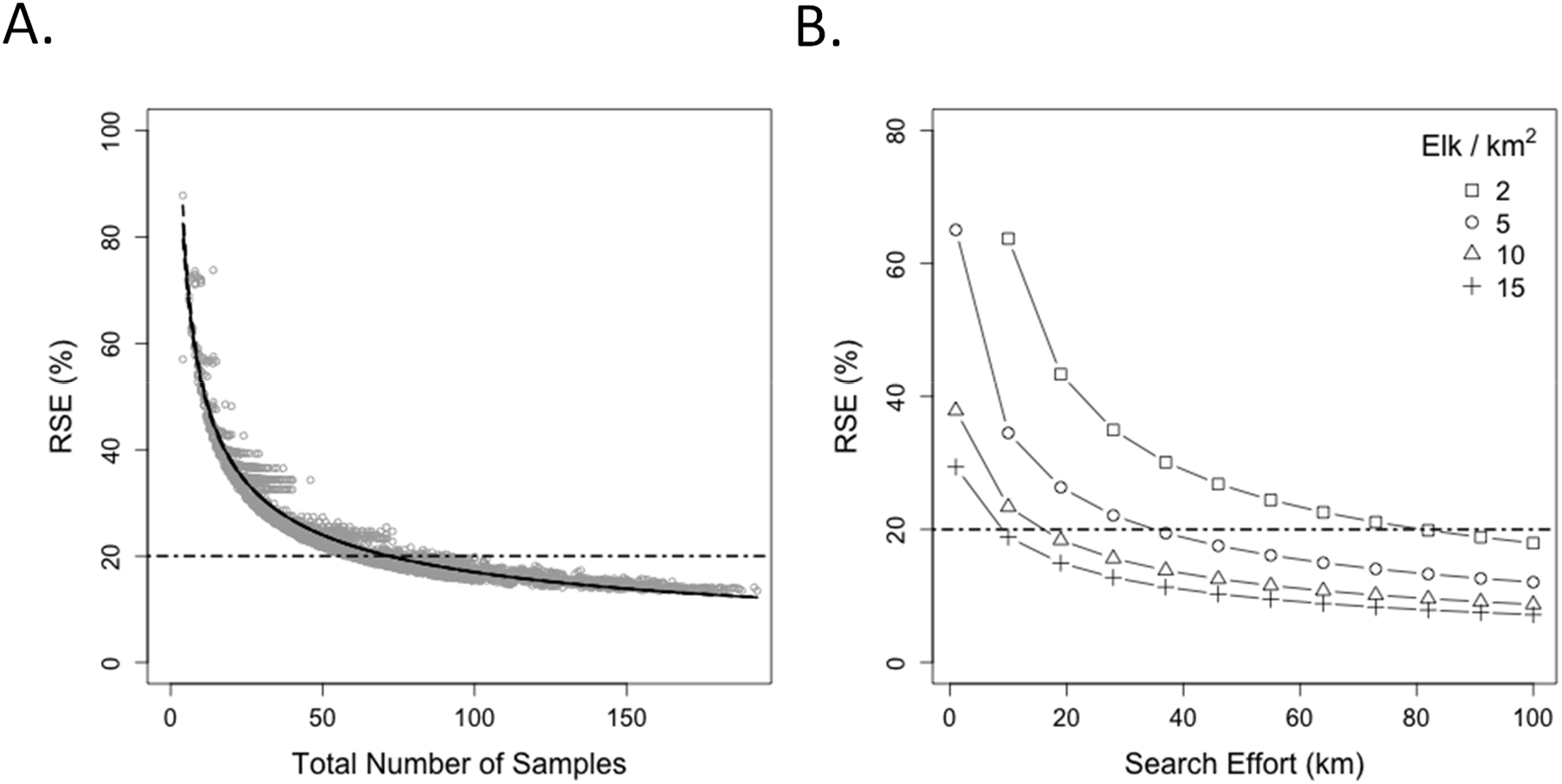
Predicted relative standard error 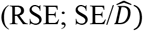 of density estimates from simulated reductions of a full dataset from a noninvasive genetic spatial capture recapture study on an enclosed population of tule elk of known density. Simulated reductions resulted in 3000 datasets that varied sampling effort and elk density by number of 250 × 250m plots (15, 20, 25, 20, and 25 plots) and number of elk (10, 20, 30, 40, 50, and 60 elk). Model predictions are from generalized linear models and are based on (A) the total number of successfully genotyped pellet samples used for each model (RSE ~ number of samples) and (B) the simulated elk density and the total kilometers of search effort within the subsampled plots for each model, including an interaction effect (RSE ~ search effort + elk density + search effort * elk density). The dashed line at 20% RSE represents a hypothetical maximum threshold for precision.

When modeling RSE by density of elk and search effort, including an interaction term resulted in the more parsimonious model, though this model was still well below the top model (ΔAIC = 1,102.52). Results from the Farrar test indicated that the two predictors were not correlated. As was expected, the RSE declined (i.e., precision improved) as the density of elk and search effort increased, with a positive interaction term (β = 0.12, P < 0.001). The model predicted that, if the true density of elk was 10 elk/km^2^, or 35 elk in the enclosure, obtaining an RSE ≤20% was feasible with <20 km of search effort in the 3.49 km^2^ enclosure. If *D* was 2 elk/km^2^, or there were approximately only 7 elk in the enclosure, it would require >80 km^2^ of search effort to reach that same level of precision (Fig. 2A).

#### Random Encounter Model

Between 20 July and 30 August 2016, we obtained an average of 18.33 (SE = 1.72) elk photos across all 18 cameras, resulting in a trapping rate (*y*/*t*) of 0.44 photos/camera/day. The recommended minimum sample size per survey to obtain results with reasonable precision is 10 photos, though a sample size of ≥20 photos is ideal (Rowcliffe et al. 2008). Bootstrapping estimated the standard error of y/*t* as 0.035. We estimated *v* to be 1.77 (SE = 0.43) km/day. Group size (g) was right-skewed, ranging from1–42 individuals, and the back-transformed mean of the natural log-transformed group counts was 3.76 (SE = 1.03) elk. Group structure was highly fluid throughout the survey, with individuals breaking off and rejoining groups, but in general, there were 2 larger cow and calf groups that included a dominant bull, and several smaller, satellite bull groups and lone bulls that often joined the larger groups temporarily. We calculated camera parameters *r* to be 12.8 m and *θ* to be 0.72 radians. Based on these parameter estimates, 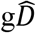 by the REM model using photos captures from all 18 cameras was 22.28 (SE = 7.86) elk groups/km^2^, with an RSE of 35%. This translated to an estimate of 77.76 (SE = 27.43) elk groups in the enclosure. Assuming an average group size of 3.76 elk, this would result in an estimate of 292.37 (SE = 130.58) elk. In sensitivity analyses for camera parameters, *r* appeared to have a much stronger negative per meter influence on 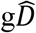 results than a per radian change in *θ* (Fig. 3A–B).

**Figure 3.**
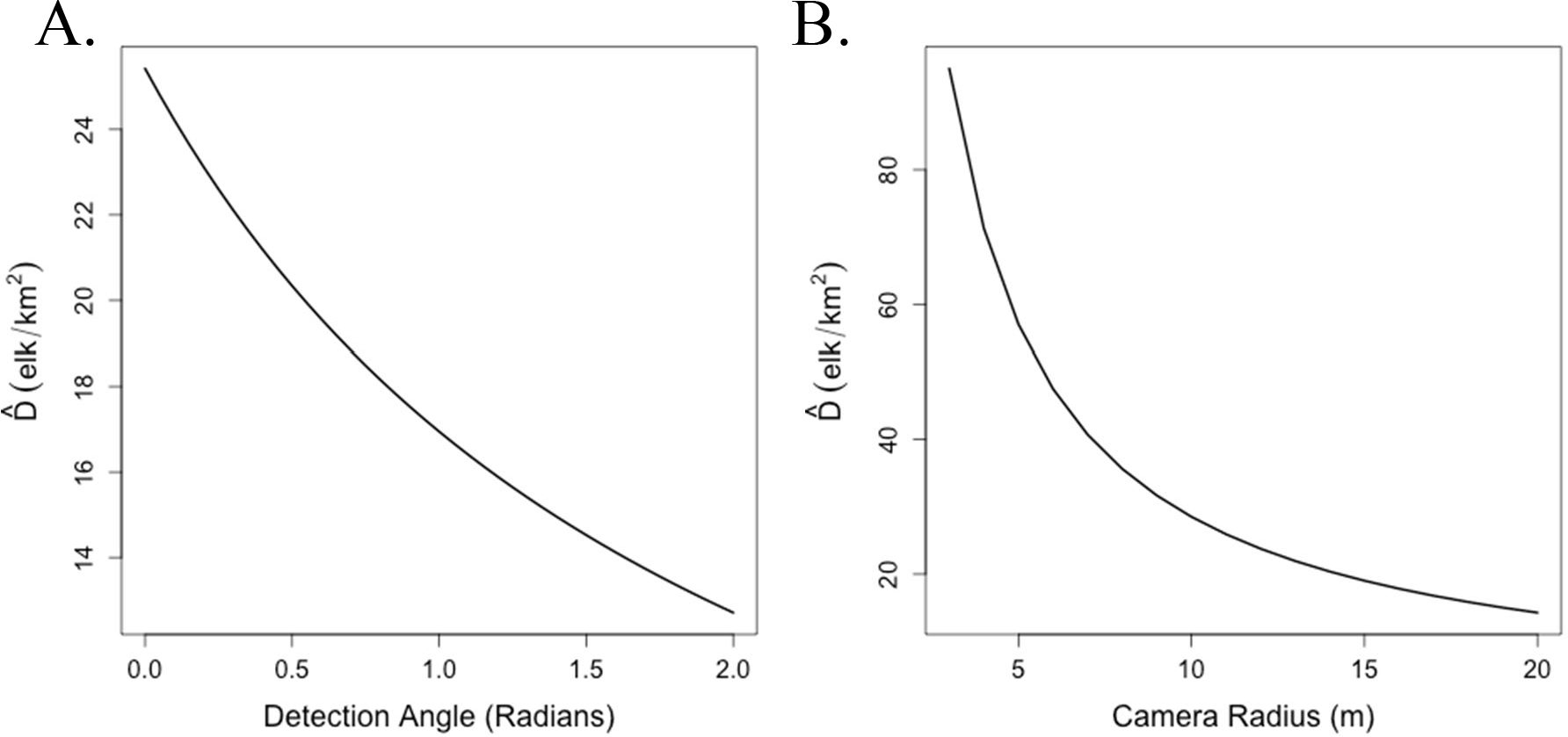
Results from sensitivity analyses that show the effect on the estimated density of elk using random encounter models (REM; Rowcliffe et al. 2008) of camera detection parameters (A) detection angle and (B) camera radius.

## Discussion

Non-invasive fecal genetic sampling and SCR models performed well in estimating the true individual density of elk, both with the full dataset and in most subsampling scenarios of reduced sampling and elk densities, despite the aggregating movement patterns of elk. In our study, the null model for detection probability was the most parsimonious model that estimated *N* with the least bias (s0), though alternative models that varied detection parameters by sex were <4 ΔAIC of the null model, and provided identical estimates of *N* (s1 and s2). Heterogeneity mixture models that allowed for different detection probabilities for 2 latent classes of individuals resulted in estimates of *N* 23–25% higher than the true *N*, even though these were ranked higher than the null model by AIC. Previous simulation studies have similarly demonstrated that using conventional measures, such as deviance or AIC, to select among individual heterogeneity models can fail to select models that provide unbiased estimates of *N* (Dorazio and Royle 2003), suggesting caution should be used in employing such models to estimate *N* or *D*.

The estimates of *N* from reduced datasets mostly had <10% relative bias compared to the census number of 72 elk within the enclosure. At the lowest levels of sampling (15 plots) and the lowest numbers of elk (10 and 20 elk), 3 models had to be eliminated because they did not provide reasonable estimates of *N* or did not provide SE estimates. While the RB in the remaining models was only −5% for both scenarios, the necessity of eliminating the replicates demonstrated that there is a minimum threshold of sampling effort required to obtain reliable estimates of *N* or *D*, especially at lower animal densities. In other words, sampling at these lower levels (15 250 × 250 m plots at densities of 2.9–5.7 elk/km^2^) will still produce relatively unbiased estimates of *N* on average, but on occasion, surveys will fail to produce reliable estimates. Conversely, the only scenarios that resulted in >10% |RB| were those at higher animal densities (>30 elk in the enclosure), when sampling only 15 plots, suggesting that sparse sampling at high densities can negatively bias estimates of *N* (Table 3).

The best predictor of precision (RSE) was the number of pellets used to construct detection histories. The number of pellets successfully genotyped and used to build encounter histories would therefore be a useful minimum sampling goal in monitoring studies using non-invasive genetic SCR methods. We found that 72 successfully genotyped pellet samples (or 1 per individual in the population) from a single season’s survey could serve as a preliminary minimum sampling goal if the desired RSE was ≤20%. However, the relationship between RSE and total number of fecal samples available for SCR modeling might change with factors affecting the detectability of individuals in the sampled population. These include deposition rates, factors affecting genotyping success rates, such as diet and the degradation rates of pellets, and possibly the scale of movement of the animals, and visual detectability of the samples themselves in different habitats (Neff 1968, Jung and Kukka 2016, Pfeffer et al. 2018). For example, in terms of scale of movement, elk in this study were restricted to an area the size of 3.49 km^2^, but wild elk populations can have summer home ranges 20–100 km^2^ or more in size (Anderson et al. 2005). If the scale of the sample collection is appropriate to the scale of the movement of the species (i.e., search effort is distributed across the landscape to capture a representative range of recapture distances within different animal home ranges), then the relationship we found might still perform well at predicting RSE. If, however, elk tend to deposit pellets in clusters, then an insufficient number of spatial captures might result for 72 samples if the survey design is scaled inappropriately. Similarly, if there is extreme heterogeneity in detectability, 72 could be biased towards highly detectable individuals and not adequately representative of the entire population. We therefore recommend that researchers or wildlife managers use results from a pilot study, if available, to establish a minimum threshold number of samples for a desired precision level, specific to their study area and population of interest. Such a model would be helpful in optimizing the efficiency of study designs for long term monitoring projects, where precision of estimates is an important factor affecting power to detect trends in population size or density.

In contrast to fecal genetic SCR, the REM models based on camera detections performed poorly at estimating density. We observed bull elk in the enclosure frequently associating temporarily with cow and calf groups, or aggregating into small bull herds, in addition to breaking off into groups of 1. It is possible that the fluidity of group membership violated model assumptions that the groups are static (Rowcliffe et al. 2008), resulting in failure to adequately estimate *gD*. A second possibility was that *gD* was not properly estimated because of the distances between individuals. Incorporating the diameter of the group distribution across space (i.e., the area occupied by a group of elk at any one time) is necessary to estimate group density based on encounter rates using the ideal gas model (Hutchinson and Waser 2007), but was not a component of the REM group density model. If the total area covered by groups of individuals across space is not accounted for, this could lead to positive bias in the estimate of group density, especially when the group diameter is large relative the space over which groups move, as was the case in our study. A third possibility was that using hourly GPS fixes to estimate animal velocity negatively biased estimates of *v* on a per day basis, and therefore inflated estimates of *D* (Pfeffer et al. 2018). Instead of using hourly GPS fixes, more recent methods enable estimation of animal velocity using the cameras themselves, which can provide estimates of velocity while reducing costs associated with collaring live animals (Rowcliffe et al. 2016, Pfeffer et al. 2018). Additional sources of possible bias in our estimate of *v* included our small sample size (n = 2) and the time period that we sampled (16 November, 2016 and 9 December, 2016), which did not overlap with the camera survey period. Our sensitivity analyses also suggested that estimates of *D* were quite sensitive to camera detection parameters, particularly the radius of camera detection. As the radius or angle of camera detection increased, the estimated density decreased. Field methods have been developed more recently to directly incorporate variation in the camera detection parameters in the model (Rowcliffe et al. 2011).

The precision of the REM estimate of density was also lower (i.e., greater error) than the estimate from SCR using the full dataset. The RSE using all 18 cameras was 35%, compared to an RSE of 12% using the full SCR dataset. However, in reduced SCR datasets, RSE ranged from 14–54%. More cameras, camera days, or the rotation of cameras during the study would increase sample size, and therefore reduce RSE from REM methods, but in a previous study, REM methods also resulted in much lower precision compared to photographic capture-recapture methods when estimating density of zebra (Zero et al. 2013). This is not necessarily surprising, as the REM attempts to estimate detection parameters without the additional information on individual identity.

We have shown that SCR methods and noninvasive genetic sampling can provide accurate and robust estimates of individual density in elk, which can help managers estimate population numbers and detect trends in population size over time. The REM models, in contrast, did not appear to perform well for estimating group density of elk in our study, had lower precision than the SCR methods, and were highly sensitive to camera detection parameters. The subsampling we performed on the full empirical dataset for SCR also simulated more realistic densities and sampling efforts of wild elk populations, and thus inform minimum sampling requirements for planning future elk population surveys. We ignored the group structure of elk here without any apparent effect on the accuracy of our estimates, but additional research in wild settings would help to improve our understanding of the effects of dependence in elk movements on these estimators. Additionally, if group density and individual density are both parameters of interest, or assessing group structure is of interest, it may be helpful in the future to develop models that directly account for clustering patterns of individuals across space.

## Acknowledgments

This project was funded by the California Department of Fish and Wildlife, Rocky Mountain Elk Foundation (RMEF project Nos. CA160559, CA170539), and the Mammalian Ecology and Conservation Unit of the Veterinary Genetics Laboratory at the University of California, Davis. T. Keldson (USFWS), C. Langner (CDFW), and J. Hobbs (CDFW) provided logistical support.

